# Cotranslational protein folding can promote the formation of correct folding intermediate

**DOI:** 10.1101/2020.05.08.084228

**Authors:** P. Tao, Y. Xiao

## Abstract

Cotranslational folding is vital for proteins to form correct structures in vivo. However, it is still unclear how a nascent chain folds at atomic resolution during the translation process. Previously, we have built a model of ribosomal exit tunnel and investigated cotranslational folding of a three-helices protein by using all-atom molecular dynamics simulations. Here we shall study the cotranslational folding of three mainly-β proteins using the same method and find that cotranslational folding can enhance helical population in most cases and reduce nonnative long-range contacts before emerging from the ribosomal exit tunnel. After exiting the tunnel, all proteins fall into local minimal states and structural ensembles in cotranslational folding are more helical than in free folding. Importantly, for GTT WW domain, one local minimal state in cotranslational folding is known as correct folding intermediate, which is not found in free folding. This result suggests that cotranslational folding may directly increase folding efficiency by accelerating sampling more than by avoiding the misfolded state, which is a mainstream viewpoint in present. In addition, our method can serve as a general scheme to study cotranslational folding process of proteins.

**Statement of Significance:** In cell, the formations of correct three-dimensional structures of proteins, namely protein folding, are essential to human health. Misfolding can lead to serious diseases such as Alzheimer’s disease and mad cow disease. As the first step of *in vivo* folding, the effect of cotranslational folding on the correct folding of proteins has been the focus of scientific research in this century. Although some experiments have shown that cotranslational folding can improve the efficiency of folding, its microscopic mechanism is not yet clear. In this paper, we study the process of cotranslational folding of three proteins by using all-atom molecular dynamics simulations, and try to reveal some aspects of the mechanism of cotranslational folding from a microscopic perspective.

## Introduction

Cotranslational folding (CTF) is the first step of protein folding in vivo, which refers to the process of translation and folding of the nascent peptide chain along the channel on the large subunit of the ribosome, that is, the ribosomal exit tunnel, from the N-terminus to the C-terminus. This kind of folding is different from the widely studied *in-vitro* free folding (FF) due to the participation of the ribosome and the order of folding(1-4). It is generally believed that the efficiency of in vivo folding is higher than that of in vitro folding(5-11), but how does CTF as the initial step of *in-vivo* folding improve the folding efficiency remains an open question. This question can be divided into two parts: (1) What are the structural features of the nascent peptide chain when it comes out of the ribosomal tunnel? (2) Are these structural features helpful for subsequent folding?

For the first question, since the length of the ribosomal tunnel is about 100Å and the diameter is 10-20Å(12, 13), during the translation process, the nascent peptide chain moves along such a narrow and long pipeline, and its conformational changes must be affected by various factors, including spatial constraints, electrostatic interaction and many more(14-17). Both theoretical and simulation studies have shown that the ribosomal exit tunnel can promote the formation of helical structure(18-20), which is also consistent with the existing experimental data(21, 22), that is, the nascent peptide chain can only form helical conformation inside the ribosomal tunnel, and more complex structures can only be found in the vestibule of the tunnel. For example, the position of the formation of a β-sheet structure is about 25 amino acids from the Peptidyl Transferase Center (PTC) of the ribosome(23). Recently, the Deutsch group and the Gunnar group also found that larger structures can be formed in the vestibule, such as the entire zinc finger protein(24), spectrin(25), and the partial transmembrane segment of the potassium channel protein(26). Since only helical conformation can be formed inside the ribosomal exit tunnel, does it mean that the ribosomes only affect the folding of helix-type proteins?

Let’s consider the second question. The nascent peptide chain is usually not completely folded when it comes out of the ribosomal exit tunnel, and it may form a structure that does not exist *in vitro* folding. For example, Holtkamp and Kokic found that the five-helix protein HemK was in a collapsed conformation just after exiting the ribosome, and then folded to a nearly native state based on this conformation(27). It should be noted that this collapsed conformation, or intermediate state, does not exist in the folding process *in vitro*(27). The same is for proteins T4 lysozyme(28) and HaloTag protein(29). These results suggest that certain structures formed by CTF can guide the folding of proteins. However, this is not the case for all proteins. For example, the folding pathways of Ig and SH3 on the ribosome are not significantly different from that *in vitro*(30, 31). Does CTF only guide the sampling of specific proteins?

In order to answer the two questions above, we have studied the CTF process of three proteins by using all-atom molecular dynamics simulation method, two of which are all-β (GTT and SH3, PDB id are 2F21(32, 33) and 1YN8, respectively) and one is mixed α/β (CI2, PDB id: 2CI2(34)). Their experimentally determined structures are shown in Figure 1. Considering that the entire simulation system will have more than 1 million atoms after adding the water molecules into the large subunit of the ribosome, it is difficult to achieve all-atom simulations on the order of microseconds with current computational power, so we still use our previous strategy to simplify the system, that is, use a simple ribosomal exit tunnel model to replace the real ribosomal tunnel(35). The construction and usage of this model will be described in detail in the Methods.

**Figure 1.**
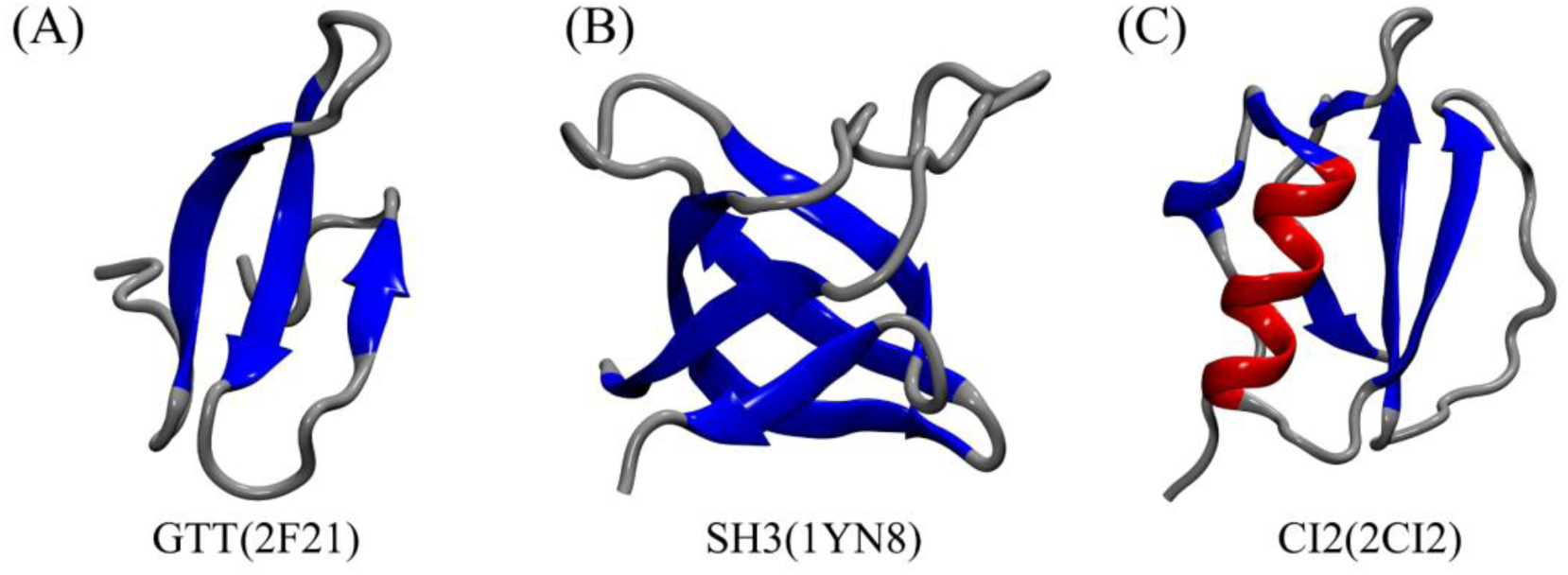
Cartoon representation of three proteins studied in this study. This figure was created by VMD program(36).

## Methods

### Construction and parameters of our simulation system

All simulations in this paper were implemented using the Amber package(37), and the protein force field used was ff14SB(38). The initial structures of the simulations were created by the tleap program(39). The dihedral angles of the main chains were set to 180°, resulting in fully extended peptides, where the N-terminus of the initial peptides were aligned to the positive direction of the x-axis. Before starting energy optimization, each protein was wrapped in a rectangular TIP3P water box, and the minimum distances between GTT, SH3 and CI2 and edges of the box were 12Å, 10Å and 10Å, respectively. Then 3 Cl^−^, 7 Na^+^ and 1 Na^+^ were added for GTT, SH3 and CI2, respectively, to neutralize the whole system. Next, two-stage energy optimization was carried out. In the first stage, the peptide was constrained by a harmonic potential of 10 kcal/mol Å^2^, and only the energy of the solvent was optimized. In the second stage, the energy of the entire system was optimized. Each stage of optimization included 2000 steepest descent optimization steps plus 2000 conjugate gradient optimization steps. After the energy optimization was completed, the system was heated from 0K to 300K in 100ps, and then maintained at 300K in another 100ps. After that, a 200ps equilibrium simulation was performed under the NPT ensemble. The time step of heating and equilibrium was 1fs and the cutoff of nonbonded interaction was 10 Å.

Using the final structure obtained from the previous equilibrium process as the initial structure, three different types of production simulations were performed for each protein, namely free folding simulation, cotranslational folding simulation at the speed of residue/2ns and residue/10ns (denoted as FF, CTF-2 and CTF-10, respectively). For each type of simulation, 8 to 10 independent trajectories were run by using different random number seeds, generating >800 μs trajectories in total (summarized in Table S1). The temperature was controlled by Langevin dynamics at 300K and the pressure was controlled by isotropic position scaling at 1bar. The time step and nonbonded interaction cutoff were set to 2fs and 9 Å, respectively. The simulated structures were saved every 10,000 steps and all simulations were performed on the GPU node of our computational cluster.

### Construction and usage of the ribosomal exit tunnel model

In order to simulate the CTF of nascent peptide chains, a rigid model of ribosomal exit tunnel was constructed, which are consisted of three parts, including a cylinder, a circular truncated cone and an infinite plane (Figure 2A). The geometric parameters of this model are shown in Figure 2B. It should be noted that the tunnel only has an effect on protein molecules, but has no effect on water molecules and ions. When an atom of the peptide reaches the tunnel wall, a completely elastic collision will occur, that is, the direction of the velocity of the atom is reversed and the magnitude is unchanged.

**Figure 2.**
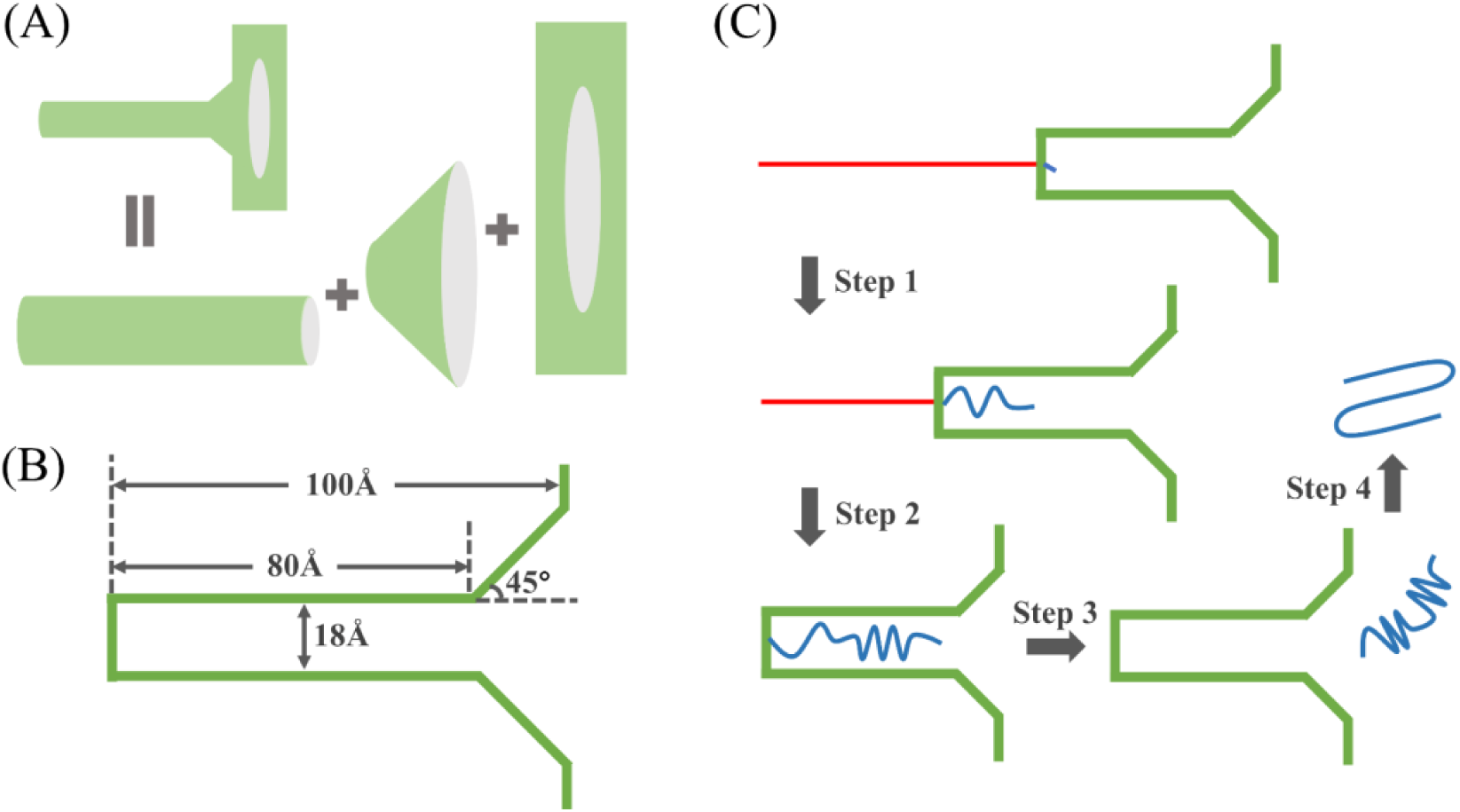
(A) Schematic diagram of the model of ribosomal exit tunnel. The green lines represent the rigid wall of the tunnel model. (B) Four geometric parameters of the tunnel model. (C) Flow chart of cotranslational folding simulation, residues with and without constraints are shown as red and blue lines, respectively.

The simulation of CTF involves four steps. The first step is to place the tunnel along the x-axis so that the peptide chain is coaxial with the tunnel (as shown in Figure 2C), then the bottom of the tunnel is moved to the minimum coordinate of the x-axis of the first residue, namely the minimum value of x coordinate of all atoms of the residue. The second to last residue of the peptide are constrained by a harmonic potential of 10 kcal/mol Å^2^ and only the first residue is allowed to move. In the second step, after a certain time, i.e., 2ns or 10ns, the bottom of the tunnel is moved to the minimum coordinate of the x-axis of the next residue, and at the same time the constraint exerted on the next residue is removed, and so on, until the bottom of the tunnel moves to the minimum coordinate of the last residue on the x-axis. The third step is to fix the position of the tunnel so that the entire nascent peptide chain can move inside the tunnel. Since the left side of the tunnel is closed, the nascent peptide chain will come out spontaneously from the right side of the tunnel after a period of time. In the fourth step, when the nascent peptide chain comes out of the tunnel, that is, all atoms of the protein molecule are on the right side of the tunnel, the tunnel model will be removed and the nascent peptide is allowed to fold freely during the rest of the simulation. The code of the tunnel model is written in Fortran language and embedded in the source code of pmemd.cuda(40, 41).

### Analysis of simulated trajectories

The calculation of the secondary structure uses the DSSP algorithm(42), which divides the secondary structure into 7 classes. In order to simplify the analysis, we further transform these 7 classes into 3 classes, among which Helix includes α-helix, 3-10 helix and π-helix, Sheet includes parallel β-sheet and antiparallel β-sheet, and Turn includes Turn and Coil. The population value of a residue is the probability that the residue is in a helix or sheet in a trajectory. It should be noted that before the peptide chain exits from the pipeline, the constrained times for the residues are inconsistent, so when comparing with FF, the population value of each residue is calculated by subtracting the corresponding constrained time. Taking the CTF of GTT at the speed of residue/10ns as an example, we take the trajectory of the first 350ns (35 residues × 10ns) to calculate the population of each residue. When comparing the population of the second residue, since the second residue is constrained for 10 ns in CTF, only the trajectory of the first 340ns (350ns-10ns) trajectory in FF is used, and so on. The calculated result using this method in FF is marked as FF-10.

In this paper, residue-residue contact is defined by the distance between Cα-atoms between two residues. When two residues are separated by more than 3 residues in sequence, and the distance between their Cα-atoms is less than 7 Å, the two residues are said to form a contact. Furthermore, if the formed contact also exists in the native structure, the contact is said to be a native contact, otherwise, it is called a nonnative contact. Before exiting from the pipeline, the calculation of the number of contacts only considers the residues without constraints. Also taking the CTF of GTT at the speed of residue/10ns as an example, when comparing the number of contacts, during the fifth 10ns (40 ∼ 50ns), since only the first 5 residues are free to fold in CTF, thus only the first five residues will be considered in FF to calculate the number of contacts, and so on.

After the peptide chain exits from the ribosomal exit channel, we compare the structural ensembles under CTF and FF and analyze the secondary structures of the two ensembles, the number of contacts, the root-mean-square-deviation (RMSD) and the radius of gyration (ROG). The calculations of RMSD and ROG exclude the loops at both ends (GTT: 7∼29; SH3: 2∼58; CI2: 5∼64, only for Cα-atoms). When clustering according to RMSD, each type of simulated trajectory are clustered into 10 classes by using the hierarchical agglomerative approach(43). The proportion of the first five classes and the RMSD values of the representative structures of these classes are summarized in Table S2. All the above analyses do not consider the first-microsecond trajectory and are done with cpptraj(44) and its parallel version cpptraj.OMP(45).

## Results

### The effect of cotranslational folding on the formation of helices depends on sequence and translation speed

First of all, we try to answer the first question: What conformation will the nascent peptide chain form in the process of CTF? What are the differences between these conformations and those formed by *in vitro* folding? Starting from the secondary structure of the nascent chain, we analyzed the proportion of each residue forming a helix and β-sheet before the nascent peptide chain completely entered into the tunnel, namely during the simulation time t∈[0, NΔt], where Δt is the translation speed and N is the residue number. Figure 3 shows the probability of each residue forming a helix at the translation speed of residue/2ns and residue/10ns. At the speed of residue/2ns, except that the helical ratio of GTT in CTF is 31% lower than that of FF, the helical ratio of SH3 and CI2 in CTF is 64% and 15% higher than in FF, respectively. At the speed of residue/10ns, the helical ratios of all three proteins in CTF are higher than those in FF, and the proportions of GTT, SH3 and CI2 increase by 161%, 100% and 41%, respectively. This shows that the influence of the ribosomal channel on the formation of the helix depends on the system and the translation speed. It is worth mentioning that the increasing rates of the helix at the speed of residue/10ns for all three proteins are higher than that of residue/2ns. Unlike the helix, the β-sheet is difficult to form, since the sampling is limited by the narrow and long space of the ribosomal exit channel, so the ratios of β-sheets of these three proteins are relatively low at both translation speeds, and there is no significant difference compared with FF (Figure 4).

**Figure 3.**
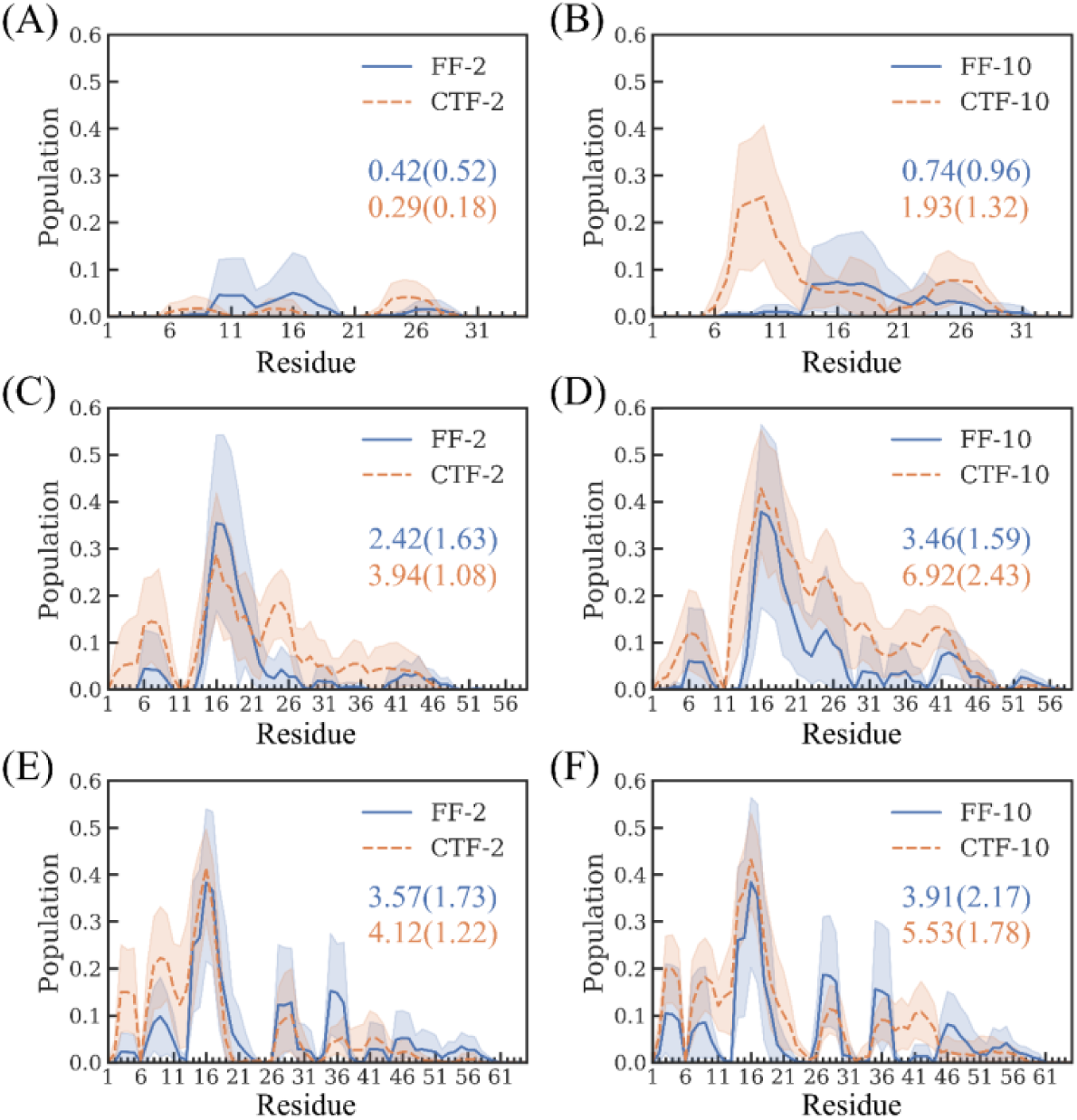
The population that each residue is in a helical structure in the cotranslational folding (orange dashed line) and free folding (blue solid line) for (A-B) GTT, (C-D) SH3 and (E-F) CI2 before exiting from the ribosomal tunnel. The corresponding shadows represent 95% confidence intervals of 8∼10 independent trajectories. The numbers outside and inside the brackets represent the average and standard deviation of the cumulative probability of all residues, respectively. The coloring scheme for numbers is the same as for lines.

**Figure 4.**
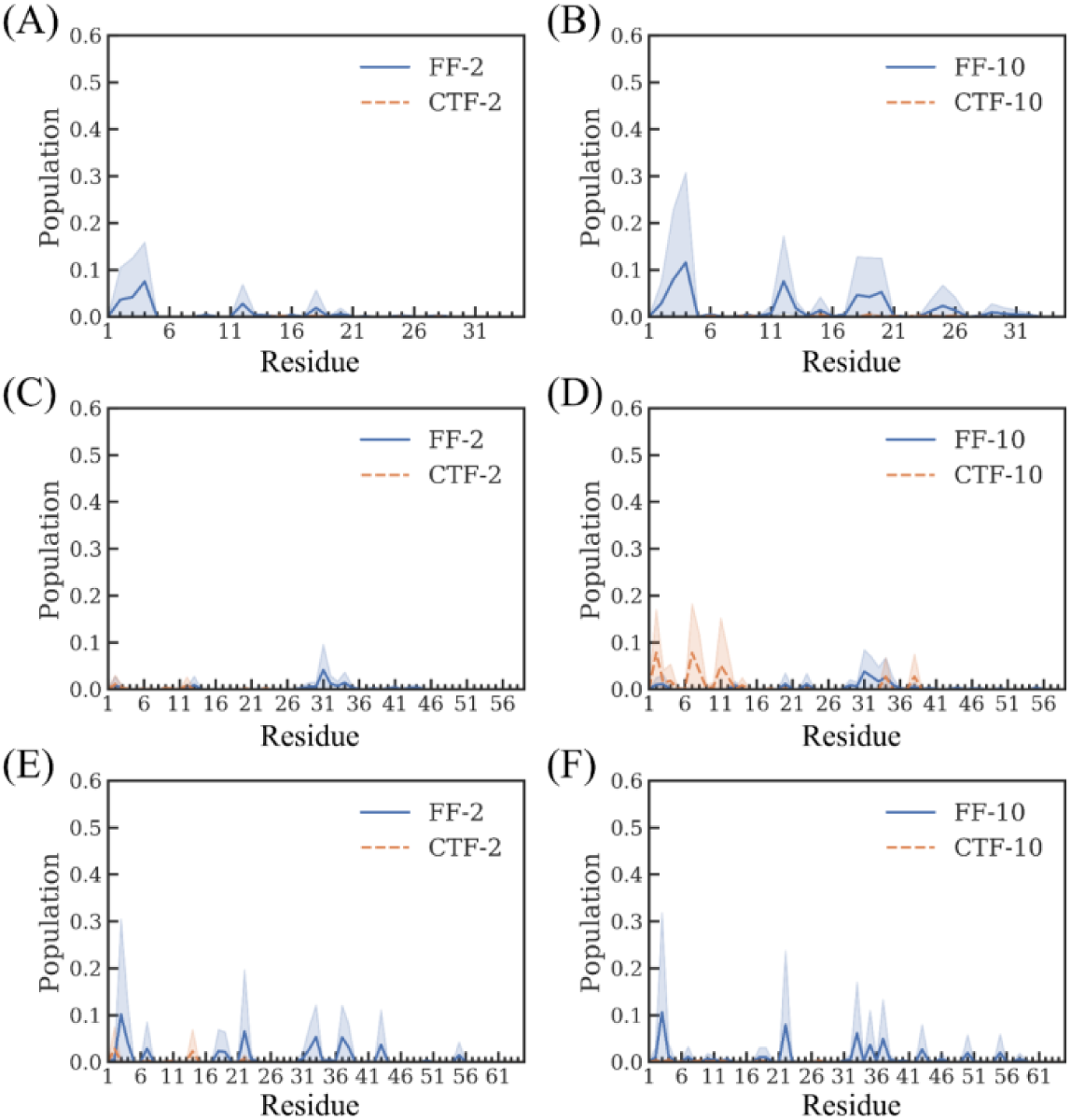
The population that each residue is in a structure of β-sheet in the cotranslational folding (orange dashed line) and free folding (blue solid line) for (A-B) GTT, (C-D) SH3 and (E-F) CI2 before exiting from the ribosomal exit tunnel.

### Cotranslational folding can significantly reduce the number of nonnative long-range contacts

In addition to the secondary structure, we also analyzed the formation of native and nonnative contacts of the nascent chain before it completely entered the tunnel, and their numbers are recorded as Q_nat_ and Q_non_, respectively. As shown in Figure 5, for the two translation speeds, Q_nat_ of these three proteins is very small (less than 5), and there is no consistent difference compared with FF. This result is in line with our expectations because these three proteins are mainly β-sheet, so native contacts are mainly long-range interactions. Surprisingly, for the two translation speeds, Q_non_ of these three proteins is significantly lower than that in FF, indicating that CTF can significantly reduce the nonnative contacts. In order to further quantify this reduction, we calculated the cumulative number of native and nonnative contacts, recorded as AQ_nat_ and AQ_non_, respectively (the area under the curve in Figure 5). As shown in Figure 6, at the speed of residue/2ns, the AQ_non_ of GTT, SH3 and CI2 are decreased to 5.8%, 29.8% and 24.7% of that in FF, respectively. Similarly, at the speed of residue/2ns, the AQ_non_ of GTT, SH3 and CI2 are decreased to 38.4%, 62.9%, and 30.0% of that in FF, respectively, which are higher than the percentages in residue/2ns.

**Figure 5.**
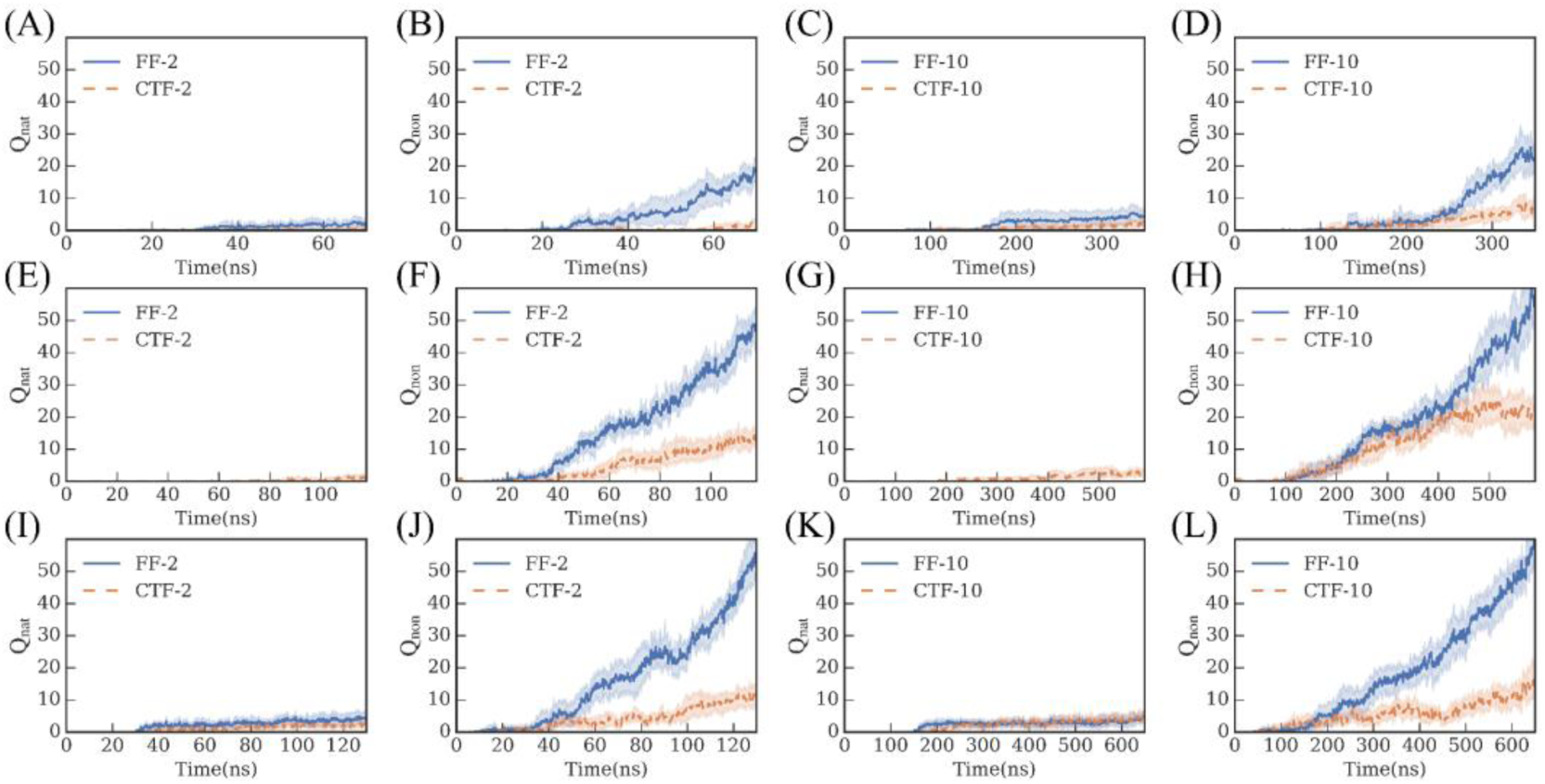
The numbers of native contacts (Q_nat_) and nonnative contacts (Q_non_) of (A-D) GTT, (E-H) SH3 and (I-L) CI2 change with time before exiting from the ribosomal exit tunnel. The meaning of the shadow is the same as in Figure 3.

**Figure 6.**
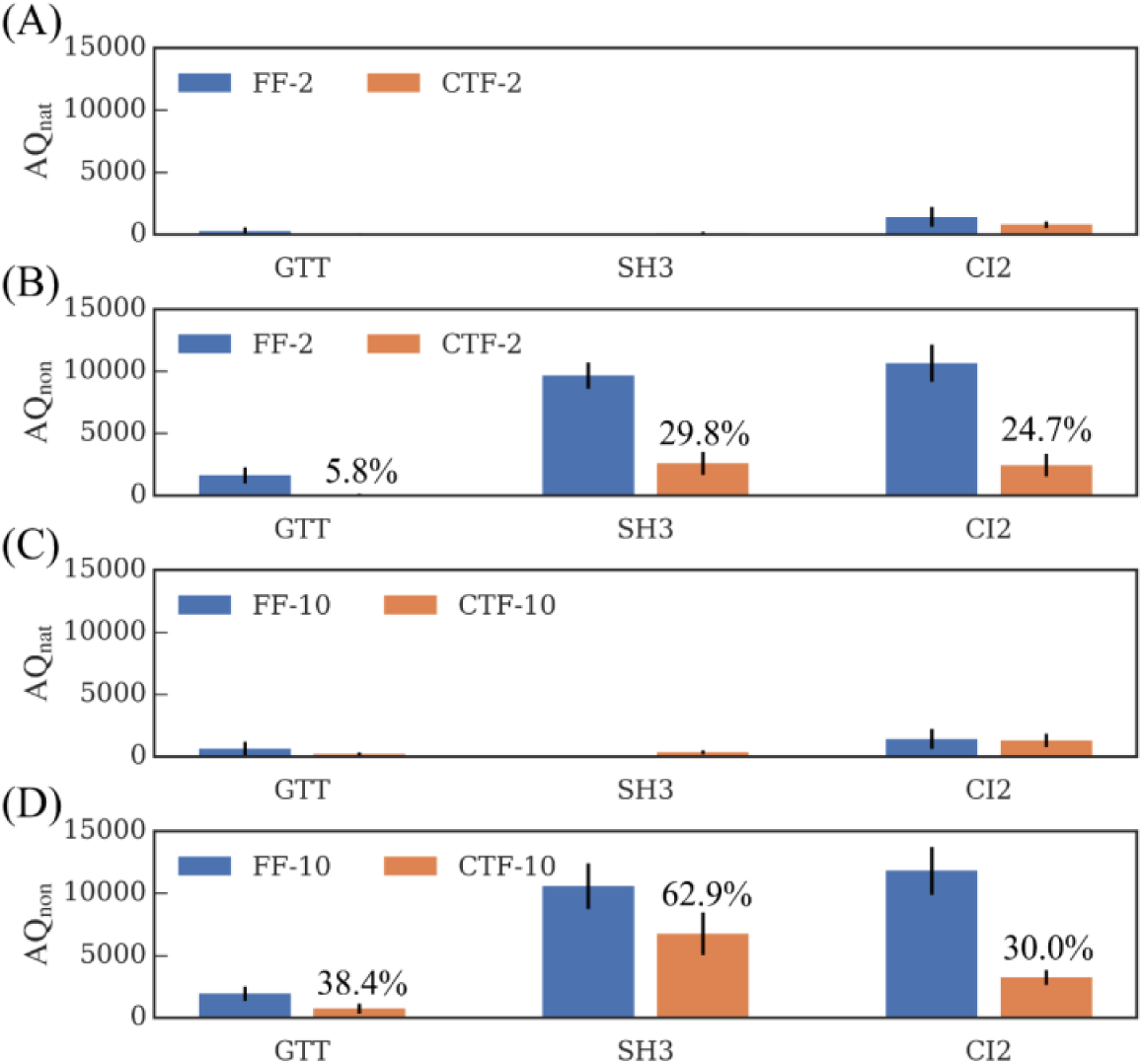
The cumulative numbers of (A, C) native contacts (AQ_nat_) and (B, D) nonnative contacts (AQ_non_) before exiting from the ribosomal exit tunnel. The labeled percentage represents the ratio of AQ_non_ in CTF to AQ_non_ in FF.

Although we already know that CTF can reduce Q_non_, maybe including long- or/and short-range contacts, it is unclear which part is reduced. Therefore, we extracted the structural ensemble of the peptide in the CTF simulation when it completely entered the tunnel and compared it with the structural ensemble that in FF at the same time (Figure 7), and analyzed the nonnative contact map of these structural ensembles. As shown in Figure 8, the contacts of the CTF ensemble are concentrated near the diagonal, which is mainly short-range contacts, while the contact of the FF ensemble includes both short-range contacts near the diagonal and a large number of long-range contacts. Therefore, mainly long-range nonnative contacts are reduced by CTF.

**Figure 7.**
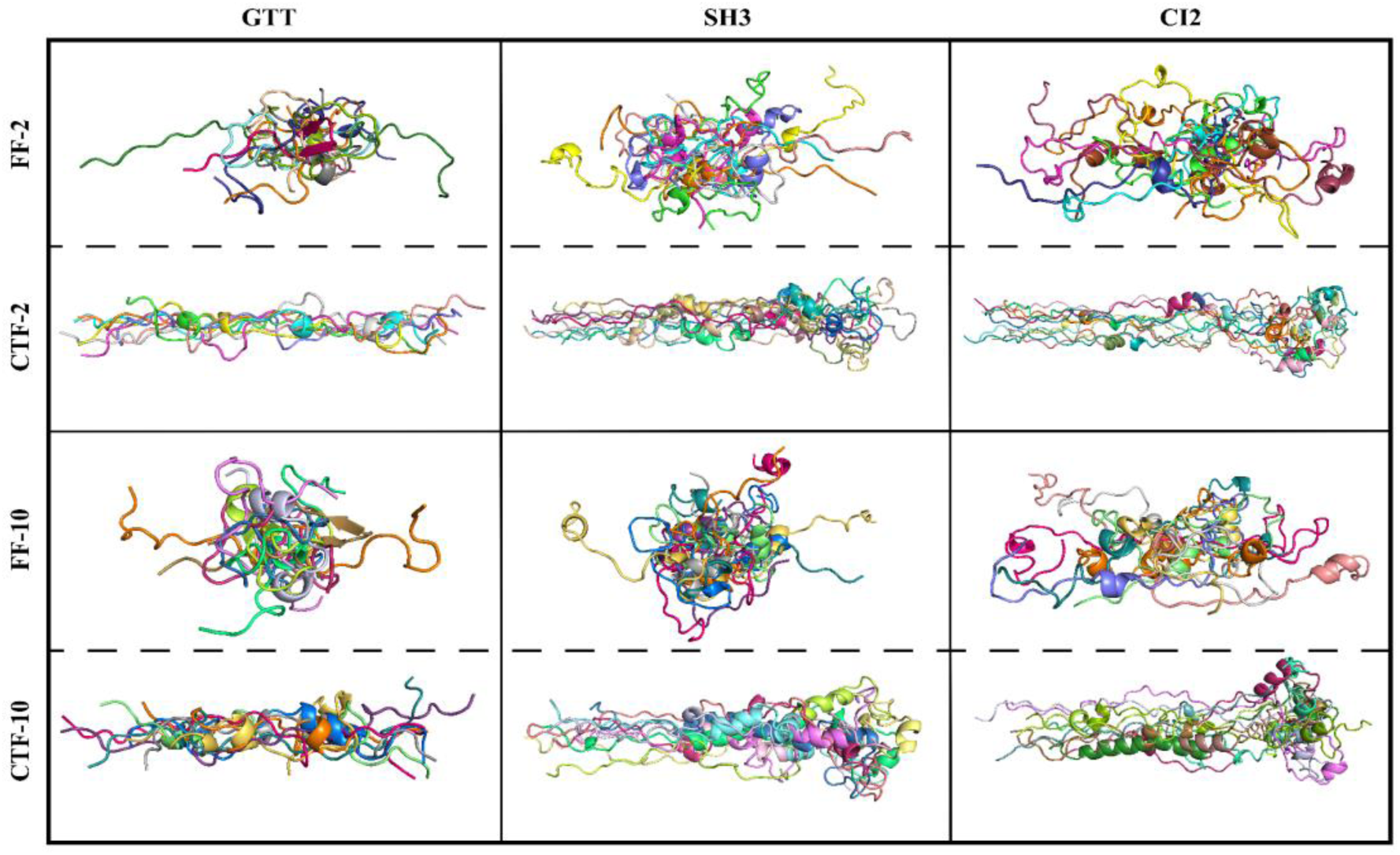
The structural ensembles of the nascent peptides in the simulations at the specific time NΔt, where Δt is the translation speed and N is the residue number.

**Figure 8.**
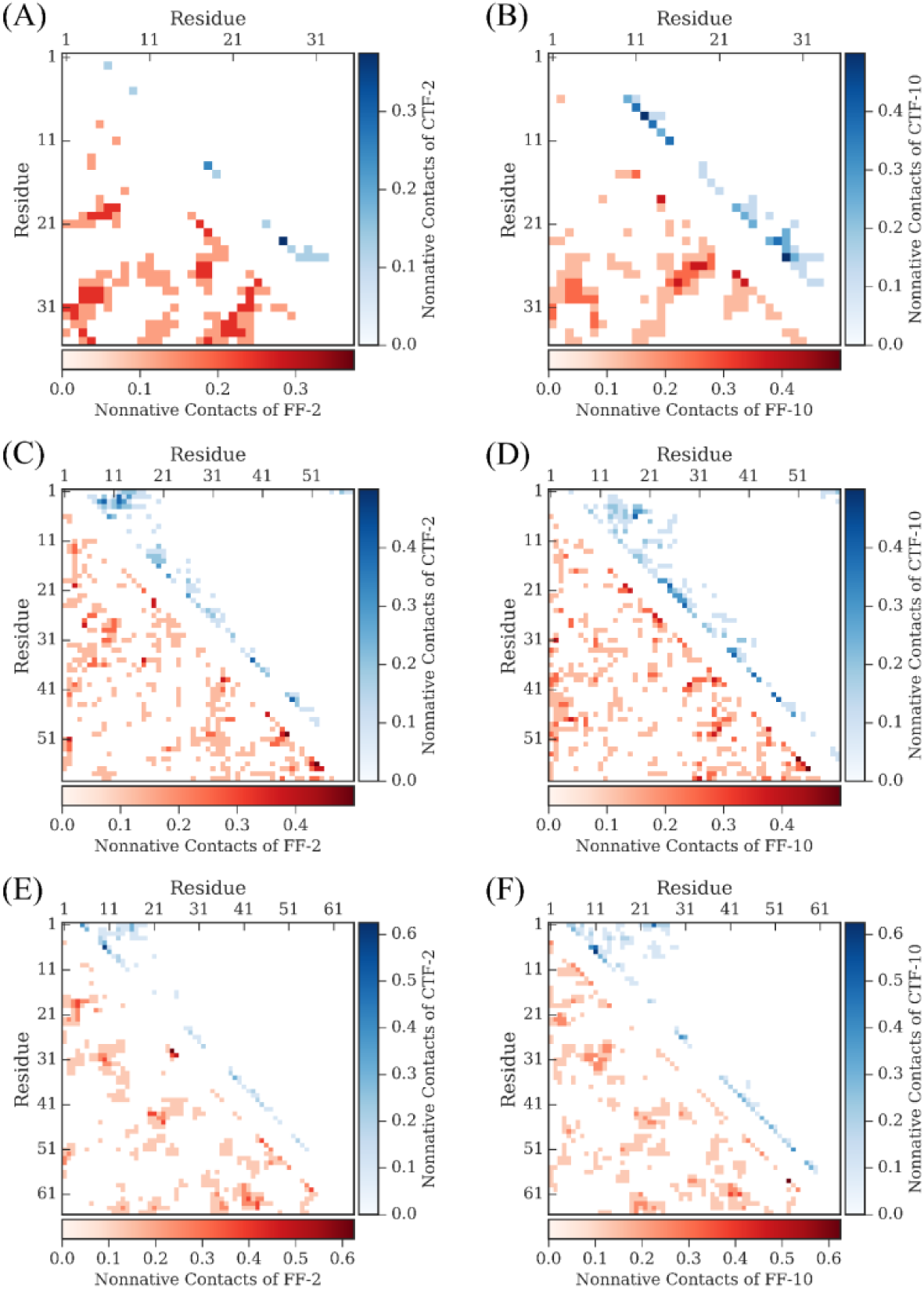
When all the residues completely enter the tunnel (that is, the simulation time equals the translation speed multiplied by the number of residues), the contact maps of the structural ensembles of (A-B) GTT, (C-D) SH3 and (E-F) CI2 under the free folding (blue, upper triangle) and co-translational folding (red, lower triangle). The darker the color, the higher the proportion of the contact in the ensemble.

### The local minimum states under cotranslational folding have a higher helix propensity

Next, turn the attention to the second question, that is to say, since CTF can make the nascent peptide chains have a higher helical propensity and fewer nonnative long-range contacts, then how do these features help subsequent folding? Due to the limited simulation time, we did not observe any complete folding events, instead, most of the trajectories fell into the local minimum state. Therefore, we can only perform statistical analysis on the trajectories of subsequent simulations, such as whether the structural features caused by the previous CTF can be maintained in the subsequent folding. To simplify the analysis, all simulations do not include the first-microsecond trajectory. First, we counted the proportion of the secondary structure of each residue in the subsequent folding of the three proteins. As shown in Figure 9, although the systems we simulated are three proteins dominated by β-sheet, the proportion of helix is much higher than that of β-sheet. In order to further quantitatively describe the difference between the helix (sheet) ratios in CTF and FF, we calculated the sum of the helix (sheet) ratios of all residues for each protein and denoted it as AP_h_ (AP_s_). Table 1 gives the average values and standard variances of AP_h_ and AP_s_ in CTF and FF. At the speed of residue/2ns, except that the helical ratio of CI2 in CTF is 7.7% lower than that of FF, the helical ratio of GTT and SH3 in CTF is 8.0% and 35.5% higher than in FF, respectively. At the speed of residue/10ns, the helical ratios of all three proteins in CTF are higher than those in FF, and the proportions of GTT, SH3 and CI2 increase by 63.4%, 56.2% and 54.0%, respectively. It suggests that the proportion of the helix that CTF raises before exiting from the tunnel can be retained to a certain extent in the subsequent folding. Thus, the local minimum state under the CTF has a higher helix propensity, especially at the speed of residue/10ns.

**Table 1.** After exiting from the ribosomal tunnel, the statistical results of the cumulative population of Helix (AP_h_) and Sheet (AP_s_), and Q_nat_ and Q_non_ of the three proteins in different simulations. The numbers outside and inside the brackets represent the average and standard deviation of corresponding quantities.

|  | GTT |  |  | SH3 |  |  | CI2 |  |  |
| --- | --- | --- | --- | --- | --- | --- | --- | --- | --- |
|  | FF | CTF-2 | CTF-10 | FF | CTF-2 | CTF-10 | FF | CTF-2 | CTF-10 |
| $AP_h$ | 4.86 | 5.25 | 7.94 | 10.93 | 14.81 | 17.07 | 10.98 | 10.13 | 16.91 |
|  | (4.20) | (3.13) | (3.19) | (4.60) | (4.92) | (5.92) | (4.94) | (3.80) | (6.31) |
| $AP_s$ | 1.38 | 2.23 | 1.33 | 2.20 | 0.85 | 0.52 | 1.11 | 3.70 | 1.26 |
|  | (1.38) | (1.79) | (2.00) | (1.57) | (0.72) | (0.62) | (0.96) | (1.34) | (0.93) |
| $Q_{nat}$ | 5.62 | 4.33 | 5.67 | 5.93 | 5.68 | 5.38 | 5.98 | 5.30 | 7.66 |
|  | (4.30) | (3.96) | (3.96) | (4.06) | (5.10) | (3.90) | (3.28) | (2.64) | (4.10) |
| $Q_{non}$ | 25.10 | 28.85 | 23.18 | 55.01 | 55.68 | 55.99 | 56.04 | 59.35 | 48.98 |
|  | (8.80) | (9.81) | (8.45) | (10.01) | (10.97) | (10.08) | (12.56) | (11.16) | (11.40) |

**Figure 9.**
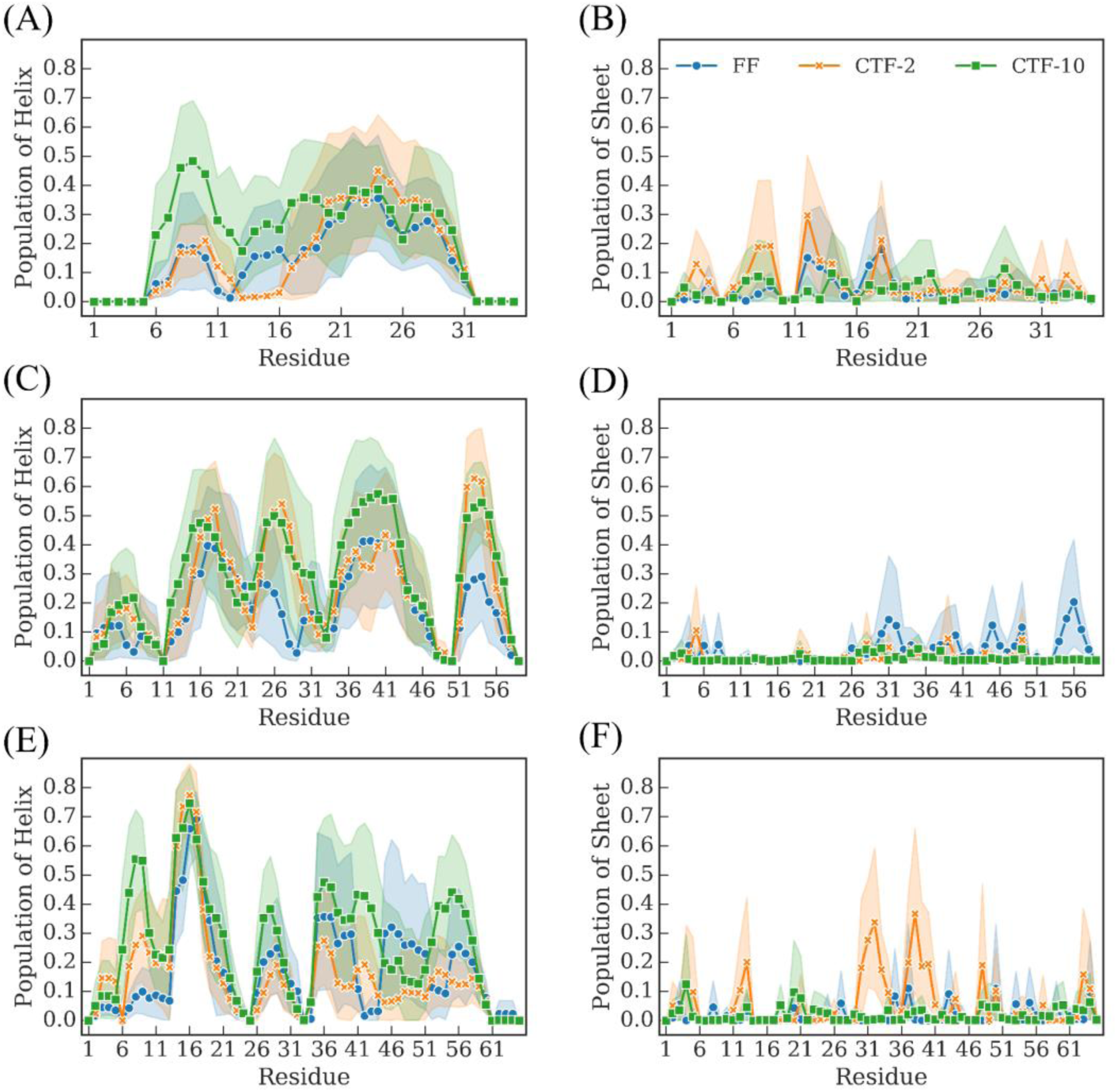
After exiting from the ribosome tunnel, the population of Helix and Sheet of each residue of (A-B) GTT, (C-D) SH3 and (E-F) CI2 under free folding (blue dots) and cotranslational folding at the speed of residue/2ns (orange crosses) and residue/10ns (green squares). The first-microsecond trajectory is not used in the analysis, and the subsequent analysis is also the same.

### Cotranslational folding can help GTT enter the correct folding intermediate

Next, we calculate the distributions of Q_nat_ and Q_non_ of the three proteins in CTF and FF. From Figure 10, we can intuitively see that these distributions are highly coincident. The average and standard variances of these distributions are given in Table 1. For GTT and CI2, the order of the average Q_nat_ is CTF-10> FF> CTF-2, and Q_non_ is CTF-2> FF> CTF-10, For SH3, the rankings of the average values of Q_nat_ and Q_non_ are FF> CTF-2> CTF-10 and CTF-10> CTF-2> FF, respectively. These results show that the advantage of CTF in the nonnative contact established before exiting from the tunnel does not maintain in the subsequent folding.

**Figure 10.**
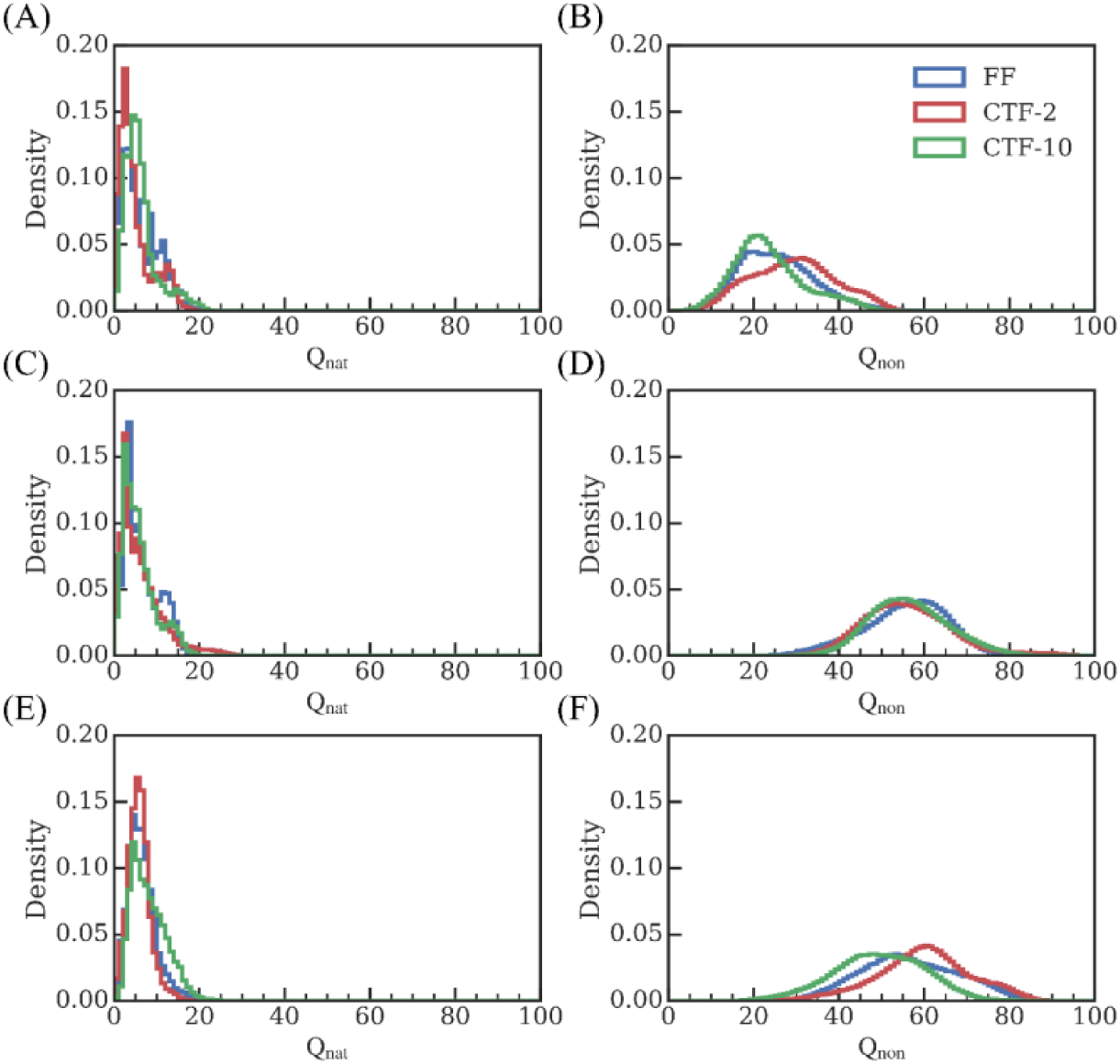
After exiting the ribosome tunnel, the distribution of Q_nat_ and Q_non_ of (A-B) GTT, (C-D) SH3 and (E-F) CI2 in free folding (blue) and cotranslational folding at the speeds of residual/2ns (red) and residual/10ns (green).

In addition to the distributions of Q_nat_ and Q_non_, we also calculated the distributions of root mean square deviation (RMSD) and radius of gyration (ROG), as shown in Figure 11. Compared with the first two distributions, the latter two distributions generally have more peaks, which means that from the first two distributions, although CTF has no obvious effect on subsequent folding, but from the latter two distributions, the local minimum states that in CTF and FF may be different. To verify this hypothesis, we clustered the subsequent trajectories according to the RMSD values and the results are shown in Figure 12 and Table S2. Taking GTT as an example, we obtained 10 classes (C1∼C10, ranking by the size of the cluster) under three types of simulations, and then compared the five largest classes. As shown in Figure 12, in FF, the first five classes do not tend to fold to the native state. In contrast, C1 in CTF-2 forms a misplaced hairpin, which may lower the barrier for the formation of the native hairpin (the first two β-sheet), while C5 in CTF-10 is in the correct intermediate state, forming a native hairpin (the last two β-sheet, Figure 13), which shows that CTF can help GTT enter the correct intermediate state.

**Figure 11.**
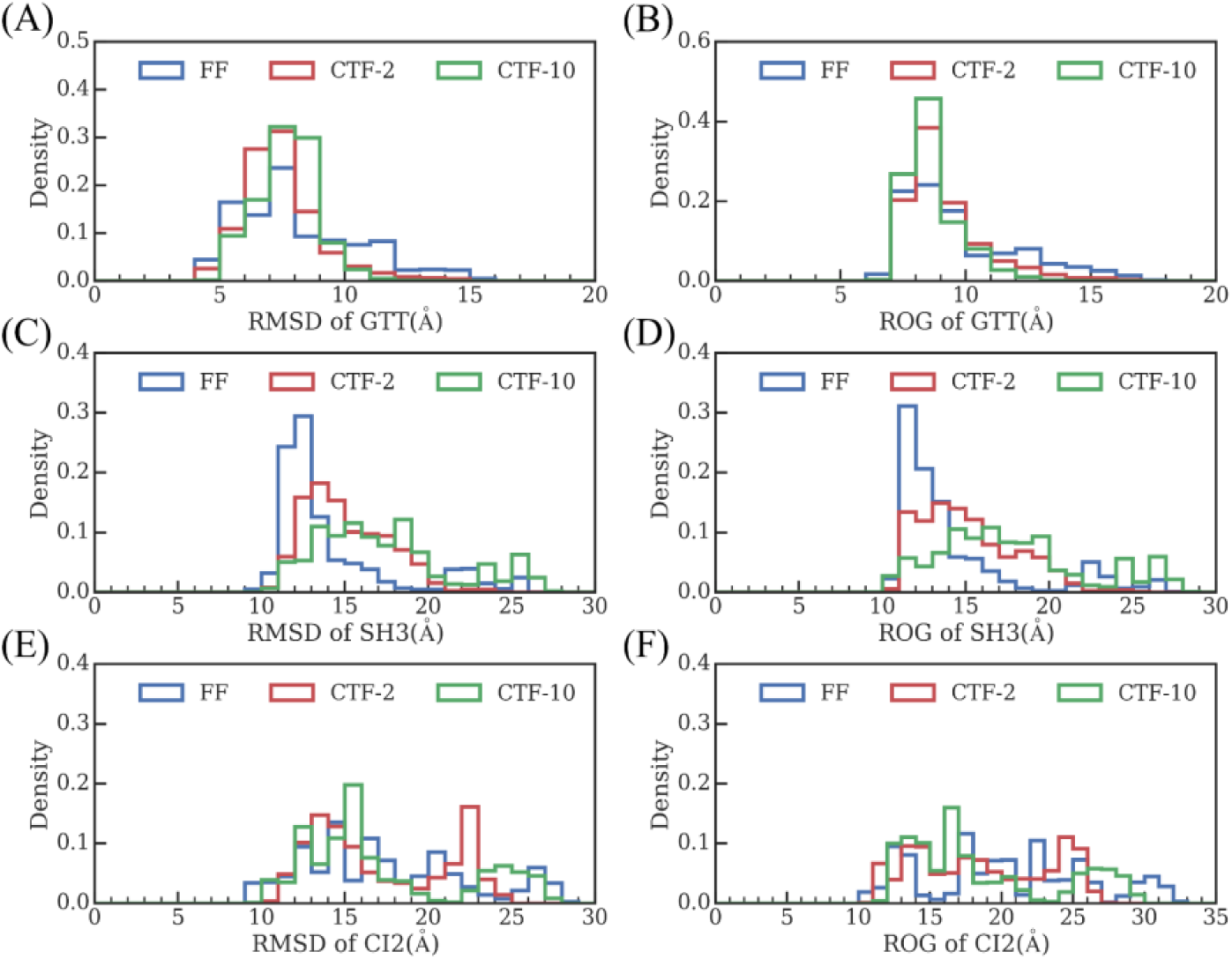
After exiting from the ribosome exit tunnel, the distribution of root-mean-square-deviation (RMSD) and radius of gyration (ROG) of (A-B) GTT, (C-D) SH3 and (E-F) CI2 in free folding (blue) and cotranslational folding at the speeds of residual/2ns (red) and residual/10ns (green).

**Figure 12.**
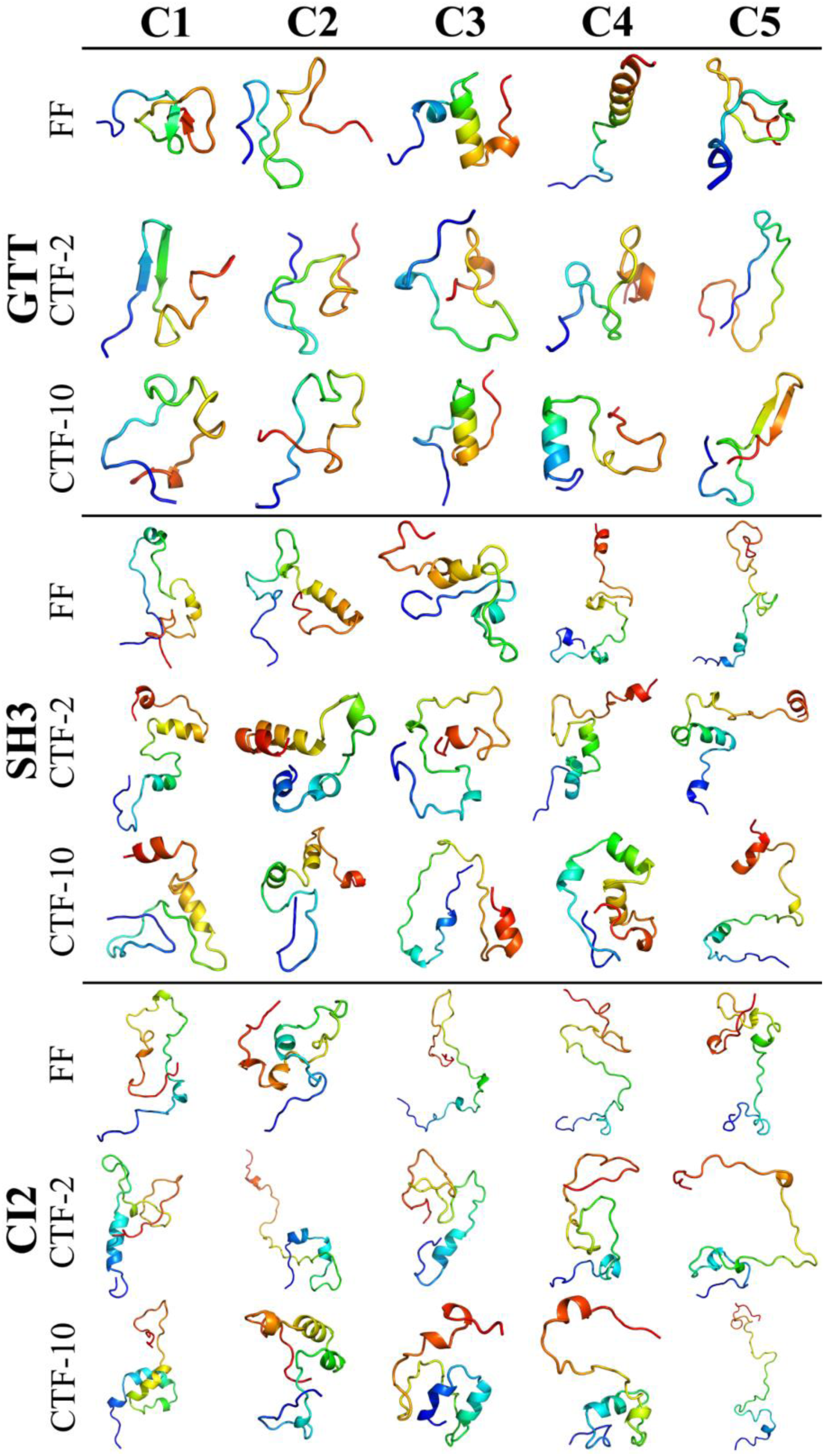
After exiting from the ribosome exit tunnel, the representative structures of the five largest clusters of three proteins in three types of simulations. The RMSD values of the representative structures and the population of clusters are summarized in Table S2. This figure was prepared with pymol(46).

**Figure 13.**
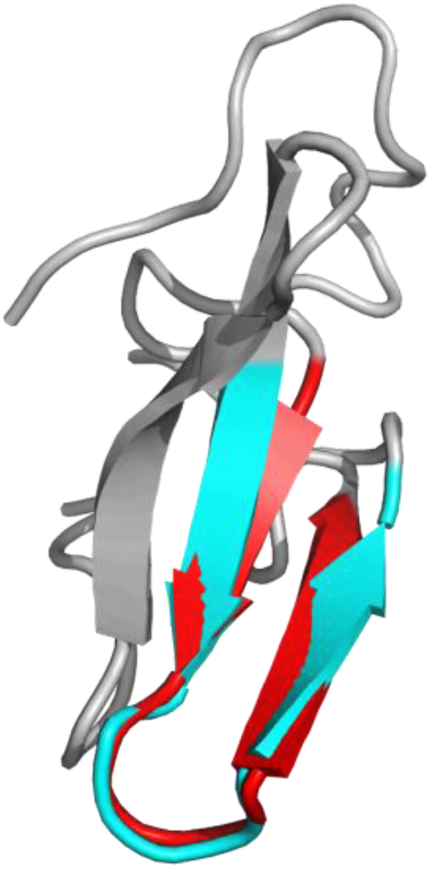
The fifth cluster C5 in CTF-10 has a native hairpin, which is the feature of on-pathway folding intermediate. The RMSD of the alignment of the hairpin is 0.99Å. Simulated and native structures are colored in red and cyan, respectively.

## Discussion

In this paper, we used all-atomic molecular dynamics simulation to study the CTF process of three proteins GTT, SH3 and CI2, and found that the helical population of the nascent peptide chain can be enhanced by CTF in most cases and lower translation speed will produce more helix. This result is easy to interpret because the nascent peptide chain has a longer time to sample the local conformation at a lower speed. In addition, CTF can also significantly reduce the number of nonnative long-range contacts. Some studies showed that, in this folding way, the nascent peptide chain can avoid falling into a misfolded state to improve folding efficiency(28, 29). This mechanism is reasonable and convincing. However, the nascent chain also rapidly collapses to local minimum states, most of them are misfolded state, in which Q_non_ is much larger than Q_nat_. Therefore, the role of CTF is beyond to prevent misfolding of nascent peptide chains. Moreover, although the distributions of Q_nat_ and Q_non_ are similar in CTF and FF, the local minimum states in CTF and FF are different. Through cluster analysis, we found that GTT is more likely to collapse to the correct intermediate state in CTF, which is not found in FF. Unfortunately, due to the limited simulation time, this phenomenon was not found for SH3 and CI2. However, for CI2, the probability of the formation of the native helix in CTF is significantly higher than that in FF, which may also be beneficial to the folding of CI2. As for SH3, we did not find that CTF has a significant effect on its subsequent folding, which is also consistent with previous research(30).

Another useful result is that CTF can significantly increase the proportion of helices and maintain a high proportion of helices in subsequent folding. So what effect does a high proportion of helices have on folding? For proteins that are mainly-α, a high proportion of helices may directly promote nascent peptide chains to form considerable native structures. For example, we recently found that CTF allows the formation of the entire N-segment of HP35 (consists of two α-helices) in the tunnel(35). Then what about mainly-β proteins? Ten years ago, Cruzeiro proposed the vibrational excited states kinetic mechanism (VES KM)(47, 48), who believed that CTF of protein could be divided into three steps. The nascent peptide forms an all helical structure in the tunnel at the first step, then the all helical structure bends at a specific amino acid site, and the third step is that the helices transform into β-sheets. According to this hypothesis, a higher ratio of helices means a better initial structure, so it will be helpful for subsequent folding. But previous experiments showed that hydrophobic sequences were easy to form helices in the tunnel, while hydrophilic sequences were inclined to form stretch conformation(49), that is, not all proteins were in a helical conformation when they exited from the tunnel, then what is the mechanism of CTF for these proteins?

It should be emphasized that the tunnel model and translation speed parameters used in this work have certain defects due to the current computing power. On the one hand, the tunnel model used in this work is rigid and regular, and the geometry of the real ribosomal tunnel is much more complicated, which may affect CTF of the nascent peptide chain, such as the position where the fold begins(50), not only that, the interaction between the ribosome tunnel and the nascent peptide can regulate the folding of the nascent peptide and even the entire translation process(14). For example, the restriction sites on the ribosomal tunnel can recognize certain sequences on the nascent peptide and cause translation pauses(51, 52). On the other hand, the actual translation speed in the cell is a few to tens of amino acids per second(53), which is about 8 orders of magnitude slower than the translation speed used in this article, which may make the sampling in the CTF more sufficient to achieve static balance sampling(54, 55). In addition, the translation speed in the cell is variable(56), and the change in translation speed may cause the nascent peptide to fold into different conformations(57, 58), and the translation speed used in this article is constant. In short, how do these defects affect the folding of the nascent peptide, and whether these effects affect the conclusions of this article need further study.

## Conclusion

In summary, a general method, combining molecular dynamics simulations and a ribosomal tunnel model, is applied to study the cotranslational folding of three mainly-β proteins. Before emerging from the ribosomal tunnel, more helical structures will be generated in cotranslation folding simulation, which will be maintained in subsequent folding. Besides, lots of long-range nonnative contacts are decreased in cotranslational folding, but the distributions of nonnative contacts in cotranslational folding and free folding are similar after the peptide exiting the tunnel. Moreover, an on-pathway folding intermediate observed in the structure ensemble of cotranslational folding does not exist in free folding, indicates that cotranslational folding can promote the formation of the correct folding intermediate. Overall, our study provides the computational evidence of the positive role of cotranslational folding on the folding efficiency.

## Supplemental Information

Supplemental Information are Figures S1-S18 and Tables S1-S2.

## Author Contributions

P. T. and Y. X. designed research. P. T. performed the research. P. T. contributed analytic tools. P. T. analyzed data. P. T. and Y. X. wrote the paper.

## Acknowledgments

This work is supported by the NSFC under Grant No. 11874162.

